# The antibody response to the glycan α-Gal correlates with COVID-19 disease symptoms

**DOI:** 10.1101/2020.07.14.201954

**Authors:** José Miguel Urra, Elisa Ferreras-Colino, Marinela Contreras, Carmen M. Cabrera, Isabel G. Fernández de Mera, Margarita Villar, Alejandro Cabezas-Cruz, Christian Gortázar, José de la Fuente

## Abstract

The coronavirus disease 19 (COVID-19) pandemic caused by severe acute respiratory syndrome coronavirus 2 (SARS-CoV-2) has affected millions of people worldwide. The characterization of the immunological mechanisms involved in disease symptomatology and protective response is important to advance in disease control and prevention. Humans evolved by losing the capacity to synthesize the glycan Galα1-3Galβ1-(3)4GlcNAc-R (α-Gal), which resulted in the development of a protective response against pathogenic viruses and other microorganisms containing this modification on membrane proteins mediated by anti-α-Gal IgM/IgG antibodies produced in response to bacterial microbiota. In addition to anti-α-Gal antibody-mediated pathogen opsonization, this glycan induces various immune mechanisms that have shown protection in animal models against infectious diseases without inflammatory responses. In this study, we hypothesized that the immune response to α-Gal may contribute to the control of COVID-19. To address this hypothesis, we characterized the antibody response to α-Gal in patients at different stages of COVID-19 and in comparison with healthy control individuals. The results showed that while the inflammatory response and the anti-SARS-CoV-2 (Spike) IgG antibody titers increased, reduction in anti-α-Gal IgE, IgM and IgG antibody titers and alteration of anti-α-Gal antibody isotype composition correlated with COVID-19 severity. The results suggested that the inhibition of the α-Gal-induced immune response may translate into more aggressive viremia and severe disease inflammatory symptoms. These results support the proposal of developing interventions such as probiotics based on commensal bacteria with α-Gal epitopes to modify the microbiota and increase the α-Gal-induced protective immune response and reduce the severity of COVID-19.

## 1. Introduction

The coronavirus disease 19 (COVID-19), a pandemic caused by severe acute respiratory syndrome coronavirus 2 (SARS-CoV-2), has rapidly evolved from an epidemic outbreak to a disease affecting the global population. SARS-CoV-2 infects human host cells by binding to the angiotensin-converting enzyme 2 (ACE2) receptor [1]. It has been established that COVID-19 mainly affects the respiratory tract, but as a systemic disease it affects multiple processes including the gastrointestinal, cardiovascular, neurological, hematopoietic and immune systems^2^. Several days after the onset of symptoms, the SARS-CoV-2 infection becomes more systemic and affecting various organs with inflammatory responses and lymphocytopenia [2]. Lymphocytopenia is likely caused by the direct lethal effect of SARS-CoV-2 on lymphocytes with the ACE2 receptor [3] and the release of pro-inflammatory cytokines such as tumor necrosis factor alpha (TNF alpha), interleukin 1 (IL-1) and IL-6 that induce apoptosis in lymphocytes [4]. The “cytokine storm syndrome (CSS)” has been associated with COVID-19 through the activation of the nuclear factor kB (NF-kB) innate immune pathway resulting in the upregulation of proinflammatory cytokines [5]. The lymphocytopenia in patients with COVID-19 along with the rise in neutrophils have been associated with a worse disease prognosis. Consequently, patients with respiratory distress syndrome in intensive care units (ICU) show lower lymphocyte counts and higher mortality when compared to other COVID-19 patients [6,7]. Additionally, COVID-19 patients suffer dysbacteriosis in the gut and lung microbiota due to enrichment of opportunistic pathogens and depletion of beneficial commensals, which suggested the development of interventions such as probiotics to reduce the severity of COVID-19 through modification of the microbiota composition [1,8,9].

Humans evolved by losing the capacity to synthesize the glycan Galα1-3Galβ1-(3)4GlcNAc-R (α-Gal), which resulted in the development of a protective response of anti-α-Gal IgM/IgG antibodies against pathogenic viruses (e.g HIV), bacteria (e.g. *Mycobacterium*) and parasites (e.g. *Plasmodium*) containing this modification on membrane proteins [10–14]. The natural IgM/IgG antibodies against α-Gal are produced in response to bacteria with this modification in the microbiota [10]. In addition to anti-α-Gal antibody-mediated pathogen opsonization, this glycan induces various immune mechanisms such as B-cell maturation, macrophage response, activation of the complement system, upregulation of proinflammatory cytokines through the Toll-like receptor 2 (TLR2)/NF-kB innate immune pathway, and TLR-mediated induction of anti-inflammatory nuclear factor-erythroid 2-related factor 2 (Nrf2) signalling pathway [14–16]. In conjunction, the immune response to α-Gal in animal models has shown protection against infectious diseases without inflammatory responses [10,12–14,17].

Based on these results, we have hypothesized that the immune response to α-Gal may play a role in the person-to-person variability in COVID-19 disease symptoms with putative protective capacity [18]. First, if the virus contains α-Gal, it would be possible to limit the zoonotic transmission of SARS-CoV-2 by antibody-mediated opsonization [18]. Secondly, boosting α-Gal-mediated protective immune and anti-inflammatory responses may contribute to the control of COVID-19 while increasing protection to pathogens with α-Gal on their surface that negatively affect the individual response to SARS-CoV-2 [14,18].

To address this hypothesis, herein we characterized the antibody response to α-Gal in patients at different stages of COVID-19 and in comparison with healthy control individuals. The results showed that while the inflammatory response and the anti-SARS-CoV-2 (Spike) IgG antibody titers increased, reduction in anti-α-Gal antibody titers and alteration of anti-α-Gal antibody isotype composition correlated with COVID-19 severity. These results suggested that the inhibition of the α-Gal-induced immune response translates into more aggressive viremia and severe disease symptoms.

## 2. Materials and Methods

### 2.1. COVID-19 patients and healthy control individuals

A retrospective case-control study was conducted in patients suffering from COVID-19 admitted to the University General Hospital of Ciudad Real (HGUCR), Spain from March 1 to April 15, 2020. The infection by SARS-CoV-2 was confirmed in all patients included in the study by the real time reverse transcriptase-polymerase chain reaction (RT-PCR) assay from Abbott Laboratories (Abbott RealTime SARS-COV-2 assay, Abbott Park, Illinois, USA) from upper respiratory tract samples after hospital admission. Clinical features, as well as laboratory determinations were obtained from patient’s medical records. The patients were grouped as (a) hospital discharge (n = 27), (b) hospitalized (n = 29) and (c) intensive care unit (ICU; n = 25) (Table 1). Patients were hospitalized for developing a moderate-severe clinical condition with radiologically demonstrated pneumonia and failure in blood oxygen saturation. Patients with acute respiratory failure who needed mechanical ventilation support were admitted to a hospital ICU. The patients were discharged from the hospital due to the clinical and radiological improvement of pneumonia caused by the SARS-CoV-2, along with the normalization of analytical parameters indicative of inflammation, such as C-reactive protein (CRP), D-Dimer and blood cell count (Table 1). Samples from asymptomatic COVID-19 cases with positive anti-SARS-CoV-2 IgG antibody titers but negative by RT-PCR (n = 10) were collected in May 22-29, 2020 and included in the analysis. Samples from healthy control individuals (n = 37) were collected prior to COVID-19 pandemic in April 2019. The use of samples and individual’s data was approved by the Ethical and Scientific Committee (University Hospital of Ciudad Real, C-352 and SESCAM C-73).

**Table 1:** Clinical parameters and laboratory tests of COVID-19 symptomatic cohort.

| Parameters | Hospital<br>discharge | Hospitalized | ICU | f-ratio | p |
| --- | --- | --- | --- | --- | --- |
| Age (years-old) | 61.0 ± 18.0 | 73.7 ± 12.6 | 57.2 ± 14.6 | 9.196 | < 0.001 |
| Neutrophils<br>(10<sup>3</sup> cells/ μl) | 7.0 ± 4.0 | 7.7 ± 4.4 | 14.2 ± 9.6 | 12.116 | < 0.001 |
| Neutrophils (%) | 68.9 ± 14.1 | 76.8 ± 10.8 | 85.1 ± 10.9 | 13.771 | < 0.001 |
| Lymphocytes<br>(10<sup>3</sup> cells/ μl) | 1.5 ± 0.5 | 1.2 ± 0.6 | 1.1 ± 0.7 | 4.223 | 0.018 |
| Lymphocytes (%) | 19.0 ± 10.1 | 13.9 ± 8.4 | 8.4 ± 7.4 | 11.521 | < 0.001 |
| Neutrophil-Lymphocyte<br>Count Ratio (NLR) | 5.4 ± 4.2 | 10.1 ± 10.0 | 18.8 ± 14.8 | 14.231 | < 0.001 |
| D-dimer (ng/ml) | 712 ± 623 | 1514 ± 1528 | 6528 ± 9436 | 7.066 | 0.002 |
| C-reactive protein<br>(CRP) (mg/dl) | 1.0 ± 1.4 | 4.4 ± 5.7 | 10.4 ± 9.7 | 17.558 | < 0.001 |

The patients were grouped as hospital discharge (n = 27), hospitalized (n = 29) and ICU (n = 25). The results (average ± S.D.) were compared between different groups by one-way ANOVA test.

### 2.2. Serum and saliva samples

Serum samples were collected for confirmed COVID-19 patients and healthy control individuals. Nursing personnel at the HGUCR extracted blood samples. Blood samples were drawn in a vacutainer tube without anticoagulant. The tube remained at rest for 15-30 min at room temperature (RT) for clotting. Subsequently, the tube was centrifuged at 1500 x g for 10 min at RT to remove the clot and obtain the serum sample. Serum samples were heat-inactivated for 30 min at 56 °C and conserved at −20 °C until used for analysis [19]. Saliva samples from asymptomatic COVID-19 cases were collected and stored at −20 °C until used for analysis.

### 2.3. Determination of antibody titers against SARS-CoV-2

Antibody titers specific for the recognition of virus infection based on IgG against SARS-CoV-2 Spike (EI 2606-9601 G) and Nucleocapsid (EI 2606-9601-2 G) proteins and IgA (EI 2606-9601 A) were determined by ELISA (Euroimmun, Lubeck, Germany) following manufacturer’s indications [19,20]. Briefly, 100 μl of the calibrator, positive and negative controls and serum samples at 1:100 dilution was added to the 96-microwell plate coated with SARS-CoV-2 proteins and incubated for 1 h at 37 °C. After washing 3 times with 300 μl/well of wash buffer, 100 μl/well of enzyme conjugate (peroxidase-labelled anti-human IgG or IgA) were added and incubated for 30 min at RT. Then, after 3 washes with 300 μl/well of wash buffer, 100 μl/well of chromogen substrate solution were added and incubated for 15 min (EI 2606-9601-2 G) or 30 min (EI 2606-9601 G; EI 2606-9601 A) at RT. Finally, the colorimetric reaction was stopped with 100 μl/well of stop solution and the absorbance was measured in a spectrophotometer (Thermo Fisher Scientific, Waltham, MA, USA) at O.D. of 450 nm. Results were evaluated semiquantitative by calculating the ratio between O.D. of the sample and the O.D of the calibrator, being those under 0.8 considered as negative and those over 1.1 as positive.

### 2.4. Determination of antibody titers against α-Gal

High absorption capacity polystyrene microtiter plates were coated with 50 ng of BSA coated with α-Gal (BSA-α-Gal, thereafter named α-Gal; Dextra, Shinfield, UK) per well in carbonate-bicarbonate buffer (Sigma-Aldrich, St. Louis, MO, USA) and used for ELISA. After an overnight incubation at 4 °C, coated plates were washed one time with 100 μl/well PBS with 0.05% Tween 20 (PBST) (Sigma-Aldrich), blocked with 100 μl/well of 1% human serum albumin (HAS) in PBST (Sigma-Aldrich) for 1 h at RT and then washed 4 times with 100 μl/well of PBST. Human serum and saliva samples were diluted 1:100 and 1:2, respectively in PBST with 1% HAS and 100 μl/well were added into the wells of the antigen-coated plates and incubated for 1 h at 37 °C. Plates were washed four times with PBST and 100 μl/well of goat anti-human immunoglobulins-peroxidase IgG (FC specific; Sigma-Aldrich), IgM (μ-chain specific; Sigma-Aldrich), IgE (ɛ-chain specific; Sigma-Aldrich), and IgA (heavy chain specific; Bio-Rad, Hercules, CA, USA) secondary antibodies diluted 1:1000, v/v in blocking solution were added and incubated for 1 h at RT. Plates were washed four times with 100 μl/well of PBST and 100 μl/well of 3,3,′5,5-tetramethylbenzidine TMB (Promega, Madison, WI, USA) were added and incubated for 20 min at RT. Finally, the reaction was stopped with 50 μl/well of 2 N H_2_SO_4_ and the O.D. was measured in a spectrophotometer at 450 nm. The average of two technical replicates per sample was used for analysis after background (coated wells incubated with PBS and secondary antibodies) subtraction. Reference values for serum immunoglobulin levels [21] were considered in the analysis of the profile of anti-α-Gal antibody isotypes.

### 2.5. Statistical analysis

The ELISA O.D. at 450 nm values were compared between different groups by one-way ANOVA test (p = 0.05; https://www.socscistatistics.com/tests/anova/default2.aspx). Pairwise comparisons between groups were conducted by Student’s t-test (p = 0.05). A Spearman Rho (*r*_*s*_) correlation analysis (p = 0.05; https://www.socscistatistics.com/tests/spearman/default2.aspx) was conducted between anti-SARS-CoV-2 Spike IgG titers and COVID-19 disease severity (2 = asymptomatic, 3 = hospital discharge, 4 = hospitalized, 5 = ICU), anti-α-Gal IgA, IgE, IgM and IgG antibody titers and disease severity (1 = healthy, 2 = asymptomatic, 3 = hospital discharge, 4 = hospitalized, 5 = ICU), and for anti-α-Gal IgA and IgG antibody titers between serum and saliva samples.

## 3. Results

### 3.1. Inflammatory biomarkers are associated with severity in COVID-19 patients

In the blood cell analysis, the ICU patients showed higher lymphocytopenia, percentage and neutrophil counts when compared to hospital discharge and hospitalized individuals (p < 0.001; Figure 1a and Table 1). The cellular and biochemical indicators of systemic inflammation, Neutrophil-Lymphocyte Count Ratio (NLR), C-reactive protein (CRP) and D-dimer levels were higher in ICU patients when compared to other patients (p < 0.002; Figure 1a and Table 1). Although more severe symptoms have been associated with elderly patients, herein older patients were recorded in the hospitalized and not the ICU group (p < 0.001; Table 1). The healthy and asymptomatic individuals did not show symptoms of inflammation. These results corroborated a higher inflammation rate in the most critical COVID-19 patients independently of the age factor.

**Figure 1:**
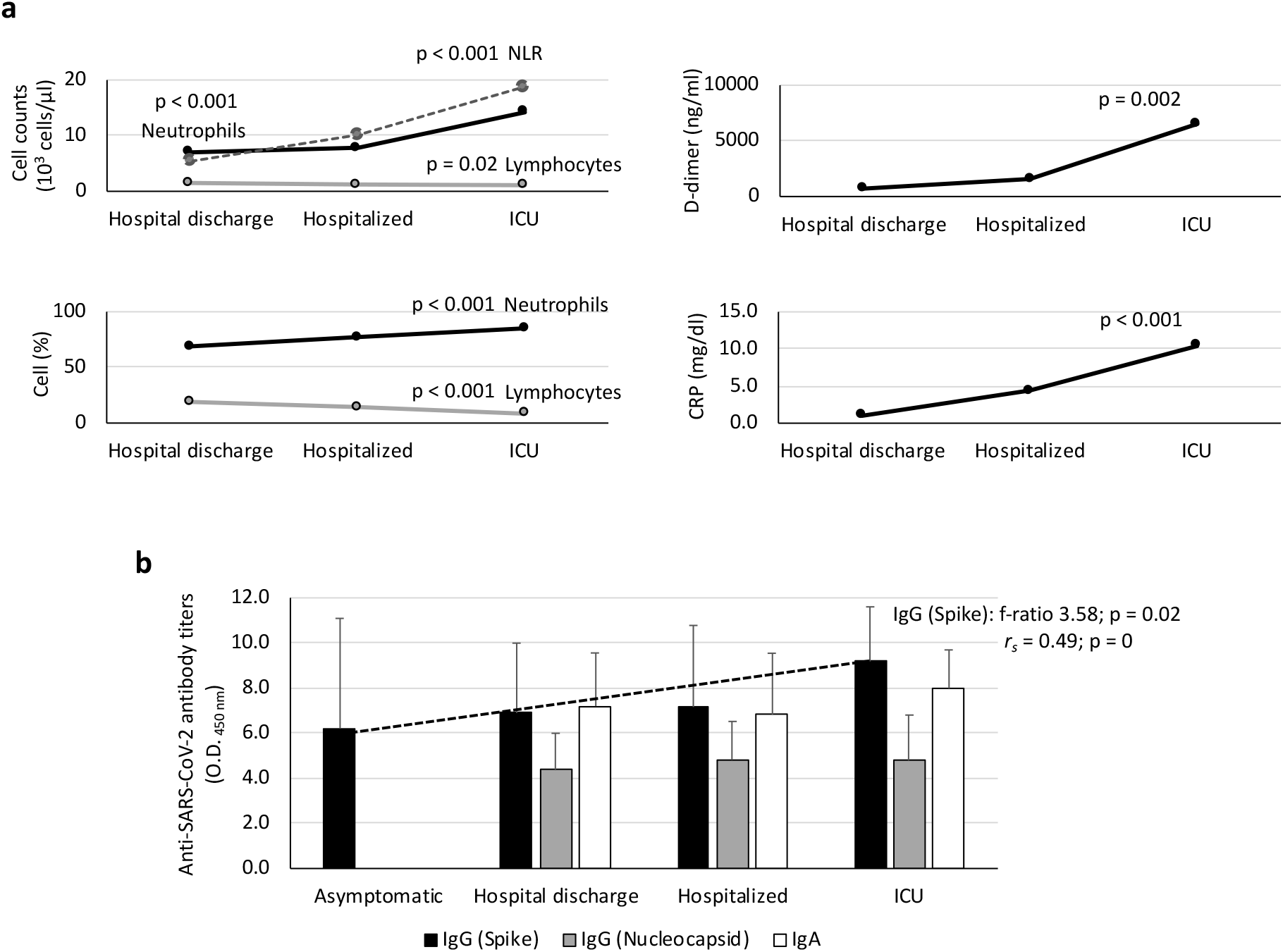
Laboratory tests in COVID-19 patients. (a) Cellular and biochemical indicators of systemic inflammation included Neutrophils (cell counts and percent), Lymphocytes (cell counts and percent), Neutrophil-Lymphocyte Count Ratio (NLR), D-dimer and C-reactive protein (CRP) levels (Table 1). (b) Serum anti-SARS-CoV-2 IgA, IgG (spike) and IgG (nucleocapsid) antibody levels were determined by ELISA. The patients were grouped as asymptomatic (n = 10), hospital discharge (n = 27), hospitalized (n = 29) and ICU (n = 25). The results were compared between different groups by one-way ANOVA test (p < 0.05). A Spearman Rho (*r*_*s*_) correlation analysis (p < 0.05) was conducted between anti-Spike IgG antibody titers and disease severity (2 = asymptomatic, 3 = hospital discharge, 4 = hospitalized, 5 = ICU).

### 3.2. Immune response to SARS-CoV-2 increased with severity in COVID-19 patients

All COVID-19 symptomatic patients showed both IgA and IgG antibody titers against SARS-CoV-2 (Figure 1b). In asymptomatic cases, only IgG antibody titers were determined, and all tested positive (Figure 1b). However, only the IgG titers against the SARS-CoV-2 Spike protein significantly increased in accordance with disease symptoms (p = 0.02; Figure 1b) with a positive correlation (*r*_*s*_ > 0; p = 0; Figure 1b). These results showed that COVID-19 patients were immunocompetent despite the inflammatory response.

### 3.3. Immune response to α-Gal varied in COVID-19 patients

The serum IgA, IgE, IgM and IgG antibody response to α-Gal was characterized in healthy individuals and COVID-19 patients at different disease stages (Figures 2 and 3a). A negative correlation was observed for IgE, IgM and IgG between anti-α-Gal antibody titers and disease severity (*r*_*s*_ < 0; p = 0; Figure 3a). The anti-α-Gal IgA antibody titers did not vary between the different groups (p = 0.21136; Fig. 3a) nor correlate with disease severity (*r*_*s*_ = 0.02; p = 0.91; Figure 3a). For anti-α-Gal IgM and IgG antibodies, the titers decreased from healthy to ICU individuals (p < 0.00001; Figures 2 and 3a). However, in asymptomatic cases the anti-α-Gal IgE titers were higher than in healthy individuals and symptomatic COVID-19 patients (p < 0.000001; Figure 3a). In COVID-19 patients, the IgE but not IgM and IgG antibody titers were higher in hospitalized patients than in hospital discharge and ICU cases (p < 0.05; Figure 2).

**Figure 2:**
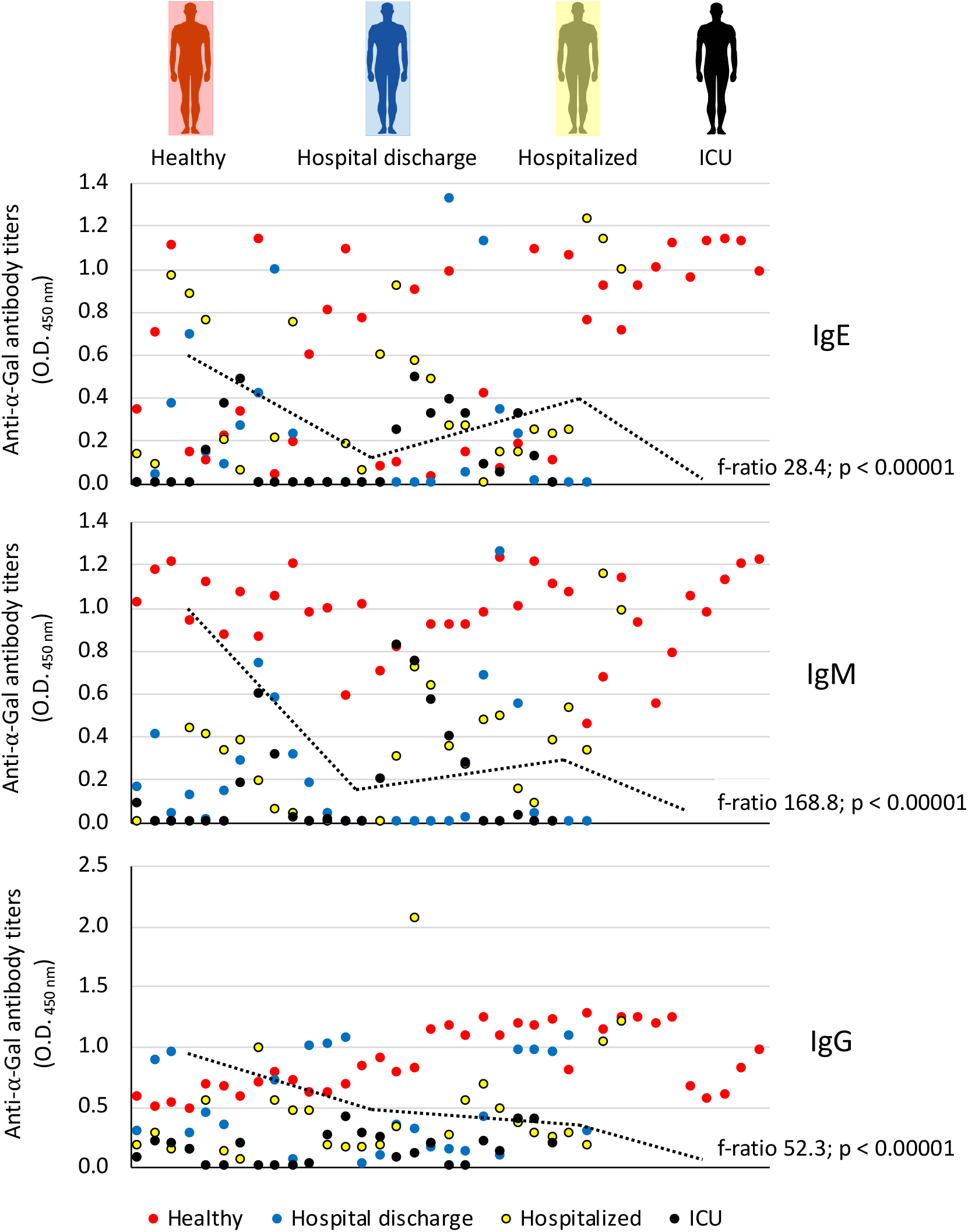
Anti-α-Gal antibody response in COVID-19 symptomatic patients and healthy controls. The IgE, IgM and IgG anti-α-Gal antibody titers were determined by ELISA. Individuals were grouped as healthy controls (n = 37), hospital discharge (n = 27), hospitalized (n = 29) and ICU (n = 25). The results were compared between different groups by one-way ANOVA test (p < 0.05).

**Figure 3:**
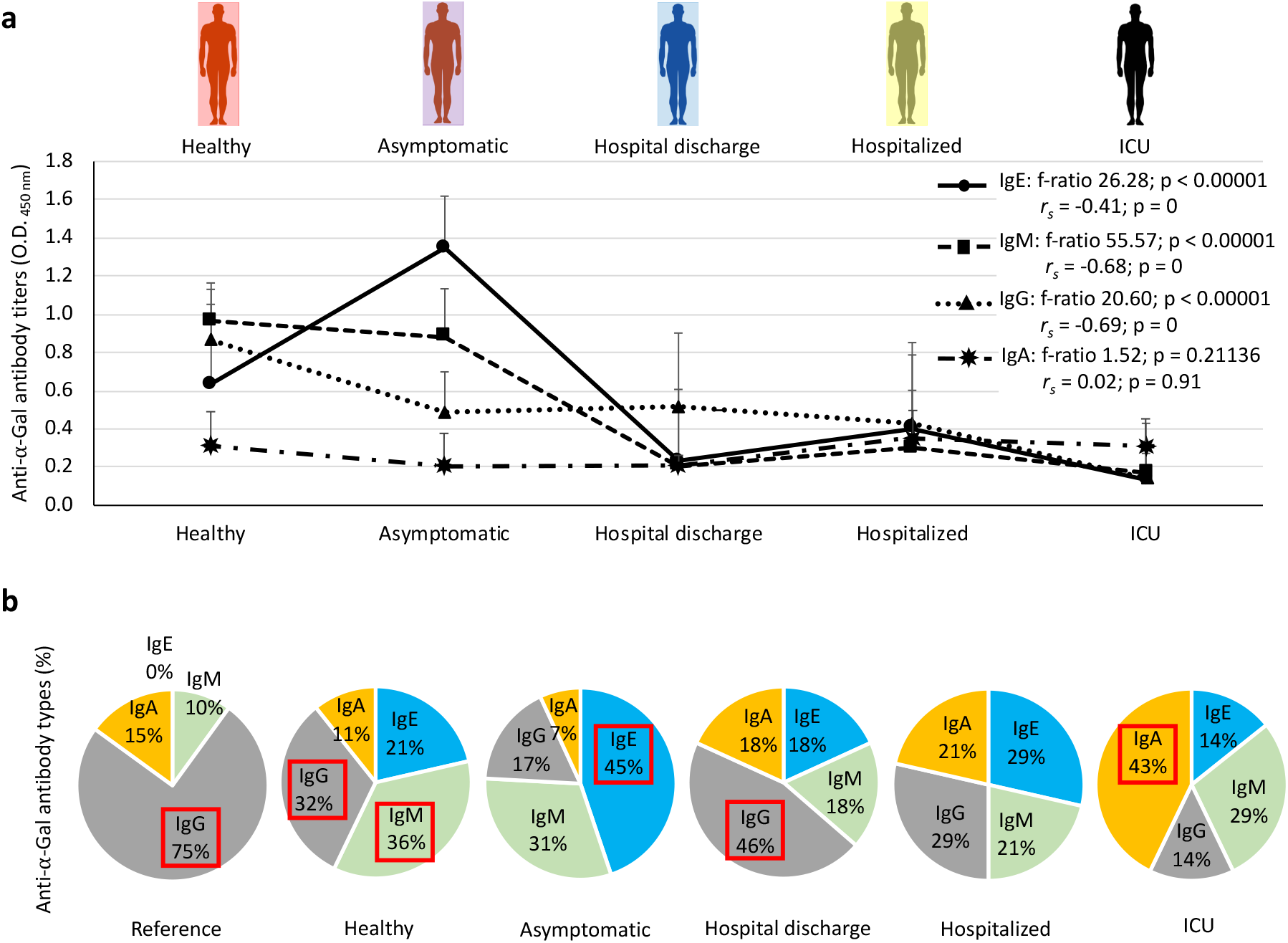
Serum anti-α-Gal antibody response in COVID-19 asymptomatic and symptomatic cases and healthy controls. (a) The IgA, IgE, IgM and IgG anti-α-Gal antibody titers were determined by ELISA. Individuals were grouped as healthy controls (n = 37), asymptomatic (n = 10), hospital discharge (n = 27), hospitalized (n = 29) and ICU (n = 25). The results were compared between different groups by one-way ANOVA test (p < 0.05). A Spearman Rho (*r*_*s*_) correlation analysis (p < 0.05) was conducted between anti-α-Gal IgA, IgE, IgM and IgG antibody titers and disease severity (1 = healthy, 2 = asymptomatic, 3 = hospital discharge, 4 = hospitalized, 5 = ICU). (b) Profile of anti-α-Gal antibody isotype (shown as percentage of antibody titers) for each group. Reference values for serum immunoglobulin levels were included. Antibody isotypes with highest representation on each group are highlighted in red.

The profile of anti-α-Gal antibody isotypes was qualitatively compared between groups including reference values for serum immunoglobulin levels (Figure 3b). The results evidenced that anti-α-Gal IgE and IgM antibodies are more abundant than reference values even in healthy individuals. However, the most abundant anti-α-Gal antibodies varied from IgM/IgG in healthy individuals to IgE (asymptomatic), IgG (hospital discharge), none (hospitalized) and IgA (ICU) in COVID-19 cases (Figure 3b).

Despite differences in absolute values due to dilutions of the samples used for ELISA (1:100 for serum vs. 1:2 for saliva), as expected, anti-α-Gal IgA but not IgG antibody titers were higher in saliva than in serum samples (p = 0.0002; Figure 4a) but without a significant correlation (p > 0.05). The saliva anti-α-Gal IgA antibody titers were similar between asymptomatic COVID-19 cases and healthy individuals (p = 0.3049; Figure 4b).

**Figure 4:**
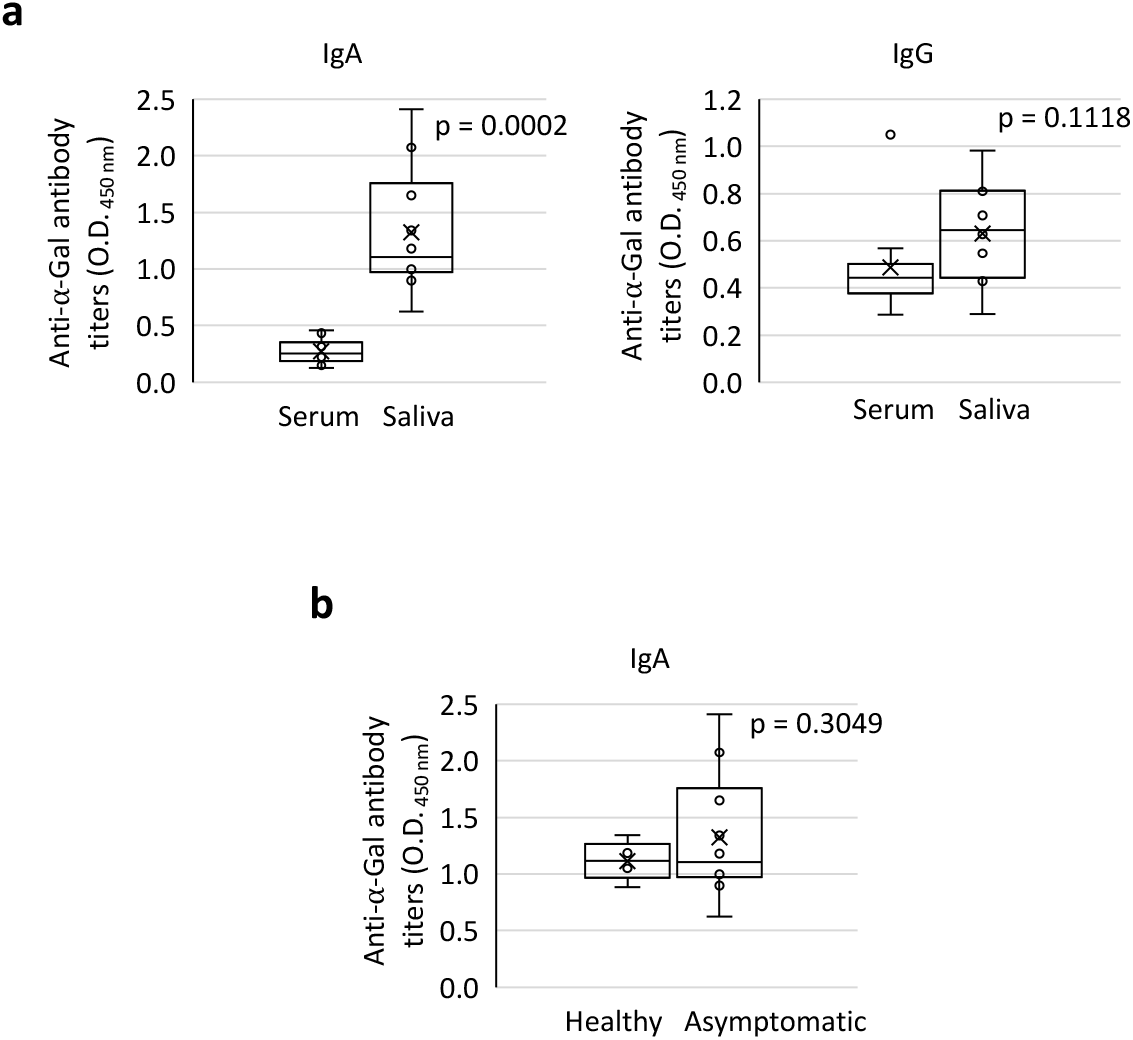
Salivary anti-α-Gal antibody response in COVID-19 asymptomatic cases and healthy controls. (a) The anti-α-Gal IgA and IgG antibody titers were determined by ELISA and compared in asymptomatic cases between serum and saliva samples by Student’s t-test (p < 0.05; n = 10). (b) The anti-α-Gal IgA antibody titers in saliva were determined by ELISA and compared between asymptomatic COVID-19 cases and healthy individuals by Student’s t-test (p < 0.05; n = 10).

## 4. Discussion

Systemic inflammation is associated with changes in quantity and composition of circulating blood cells and has been identified as the primary basic mechanism resulting in disability and increased mortality in COVID-19 [22]. As previously reported [23,24], in the blood cell analysis of cellular and biochemical indicators of systemic inflammation, the ICU patients showed higher lymphocytopenia independently of the age factor associated with more severe COVID-19 symptoms [25].

The results of our study showed a negative correlation between anti-α-Gal antibody titers and COVID-19 disease severity. However, these results raised the question of whether the observed reduction in the anti-α-Gal antibody response at the population level is a consequence or a cause of COVID-19 symptomatology. Considering that COVID-19 patients were immunocompetent with a positive correlation between anti-SARS-CoV-2 Spike antibody titers and disease severity [26], our results suggested that the decrease in the anti-α-Gal antibody response occurred by mechanisms different from humoral immunosuppression and as a consequence of SARS-CoV-2 infection.

In addition to the observed negative correlation between anti-α-Gal IgE, IgM and IgG antibody titers and COVID-19 disease severity, our results showed differences in the profile of anti-α-Gal antibody isotypes in COVID-19 cases that may be associated with different disease stages (Figure 5). These results suggested that higher anti-α-Gal IgE levels in asymptomatic cases may reflect an allergic response mediated by this glycan, which reflects the trade-off associated with the immune response to α-Gal that benefit humans by providing immunity to pathogen infection while increasing the risk of developing allergic reactions to this molecule [12,13,17]. In healthy individuals as in hospital discharge cases, the higher representation of anti-α-Gal IgM and/or IgG antibodies may be associated with a protective response to COVID-19. However, in hospitalized patients the representation of anti-α-Gal antibody isotypes did not vary, which could reflect the absence of protection. Finally, the higher representation of anti-α-Gal IgA antibodies in ICU patients may be associated with the inflammatory response observed in these cases. In accordance with these results, it was recently shown in endogenous α-Gal-negative turkeys that treatment with probiotic bacteria with high α-Gal content results in protection against aspergillosis through reduction by still unknown mechanisms in the pro-inflammatory anti-α-Gal IgA response in the lungs [27].

**Figure 5:**
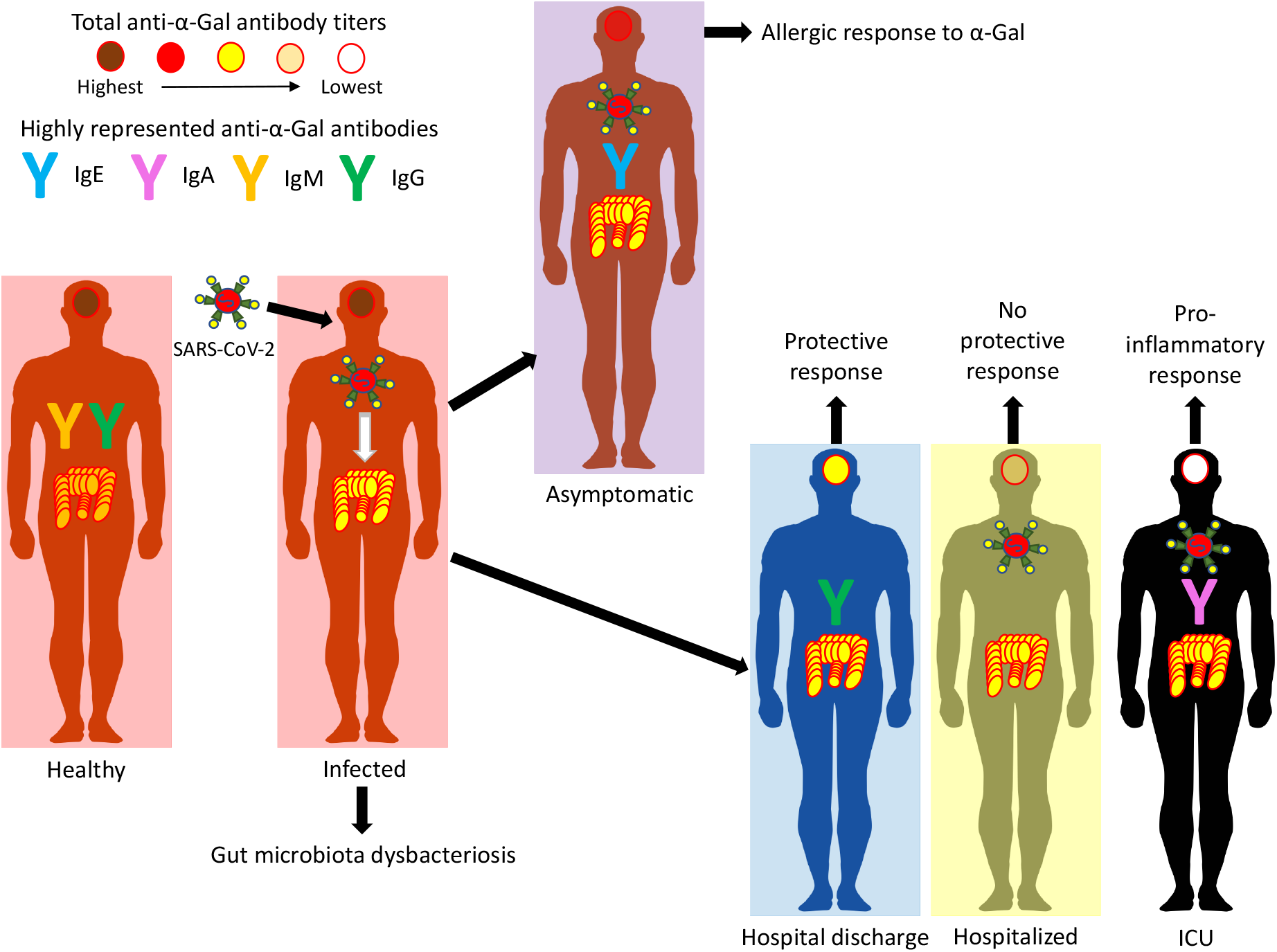
Proposed functional correlation between the anti-α-Gal antibody response and COVID-19. A negative correlation between anti-α-Gal antibody titers and COVID-19 disease severity and differences in the profile of anti-α-Gal antibody isotypes may be associated with different disease stages. Our hypothesis is that the dysbacteriosis observed in COVID-19 patients translates into a reduction in total anti-α-Gal antibody titers and alteration of anti-α-Gal antibody isotype composition due to the reduction in the microbiota of α-Gal-containing commensal bacteria.

Based on the fact that natural antibodies against α-Gal are produced in response to bacteria with this modification in the microbiota [10], our hypothesis is that the dysbacteriosis observed in COVID-19 patients [28] translates into a reduction in total anti-α-Gal antibody titers and alteration of anti-α-Gal antibody isotype composition due to the reduction in the microbiota of α-Gal-containing commensal bacteria and other still uncharacterized mechanisms (Figure 5).

Alternatively, individuals with higher α-Gal content in the microbiota may be less susceptible to COVID-19. Additionally, the pulmonary microbiota can be affected with the presence of gut bacteria in the lungs [9].

The protective response of anti-α-Gal IgM/IgG antibodies against pathogenic organisms containing this modification on membrane proteins has been well documented [10–14,17,29]. In contrast, IgE antibody response against α-Gal has been associated with the allergy to mammalian meat or alpha-Gal syndrome and other diseases such as atopy, coronary artery disease and atherosclerosis [30–33].

In preliminary analyses, it was suggested that blood type O individuals are less susceptible to COVID-19 than other blood type groups [34,35], a finding that was recently confirmed by genetic analyses [36]. ABO blood groups contain highly fucosylated antigens [37,38], a property shared with the glycans present in SARS-CoV-2^39^. For example, glycosylation in Spike asparagine (N343) is highly fucosylated with 98% of detected glycans bearing fucose residues [39]. Accordingly, the monoclonal antibody S309 that neutralizes SARS-CoV-2 binds core fucose moieties in N343 and *N*-acetylglucosamine (GlcNAc), a structural glycan found in both SARS-CoV-2 [39] and ABO blood groups [37,38]. These findings prompted us to consider that blood type O individuals could produce antibodies against A and B antigens that in addition to IgM/IgG antibodies against α-Gal, which cross-react with the structurally similar blood B antigen [40], could be involved in a polyvalent recognition of the SARS-CoV-2 Spike that may be implicated in the human protection to COVID-19.

In conclusion, according to these results and previous findings in retrovirus [41,42], the inhibition of the α-Gal-induced immune response may translate into more aggressive viremia and severe disease inflammatory symptoms [43]. These results further encourage addressing the proposal of developing interventions such as probiotics based on commensal bacteria with α-Gal epitopes to modify the microbiota and increase the α-Gal-induced protective immune response and reduce the severity of COVID-19 [8,18].

## Ethical Approval

The use of samples and individual’s data was approved by the Ethical and Scientific Committee (University Hospital of Ciudad Real, C-352 and SESCAM C-73).

## Conflicts of Interest

The authors declare that there is no conflict of interest regarding the publication of this paper.

## Author’s Contributions

J.M.U, C.G. and J.F. designed the study; E.F., M.C., C.M.C., I.G.F., J.M.U, C.G. and J.F. performed the experiments; J.F., J.M.U, E.F., M.C., C.M.C., I.G.F. and A.C-C analyzed data; J.M.U., C.G., A.M., M.V. and J.F. supervised the project. All authors wrote the manuscript.

## Acknowledgments

This work was partially supported by the Consejería de Educación, Cultura y Deportes, JCCM, Spain, project CCM17-PIC-036 (SBPLY/17/180501/000185). We thank Antonio Mas (University of Castilla La Mancha, UCLM, Spain) for the critical reading of the manuscript. We acknowledge UCLM, Spain support to Grupo SaBio. MC was funded by the Ministerio de Ciencia, Innovación y Universidades, Spain (grant FJC-2018-038277-I). IGFM was supported by the UCLM. MV was supported by the UCLM and the Fondo Europeo de Desarrollo Regional, FEDER, EU.

## Notes

### Competing Interest Statement

The authors have declared no competing interest.

